# Quantifying the Human Cost of Global Warming

**DOI:** 10.1101/2022.06.07.495131

**Authors:** Timothy M. Lenton, Chi Xu, Jesse F. Abrams, Ashish Ghadiali, Sina Loriani, Boris Sakschewski, Caroline Zimm, Kristie L. Ebi, Robert R. Dunn, Jens-Christian Svenning, Marten Scheffer

## Abstract

The costs of climate change are often estimated in monetary terms^1,2^ but this raises ethical issues^3^. Here we express them in terms of numbers of people left outside the ‘human climate niche’^4^ – defined as the historically highly-conserved distribution of relative human population density with respect to mean annual temperature (MAT). We show that climate change has already put ∼8% of people (>500 million) outside this niche. By end-of-century (2080-2100), current policies leading to around 2.7 °C global warming^5-9^ could leave one third (21-42% or 2-4 billion) of a future 9.5 billion population outside the niche. Limiting global warming to 1.5 °C could halve this exposure, reducing it by ∼1.5 billion people. For the two countries with the most people affected – India and Nigeria – reducing global warming from 2.7 to 1.5 °C results in a >6-fold decrease in the population exposed to unprecedented temperatures, MAT ≥29 °C. The lifetime emissions of ∼3.5 global average citizens today (or ∼1.2 average US citizens) expose 1 future person to MAT ≥29 °C by end-of-century. That person comes from a place where emissions today are around half of the global average. These results highlight the need for more decisive policy action to limit the human costs and inequities of climate change.

## Introduction

The COP26 meeting in November 2021 led to increased pledges and targets to tackle climate change. Yet current policies (if enacted) still leave the world on course for around 2.7 °C end-of-century global warming^5-9^ above pre-industrial levels – far from the aim of the Paris Agreement to limit global warming to 1.5 °C. Even fully implementing all 2030 Nationally Determined Contributions (NDCs), long-term pledges, and net zero targets, nearly 2 °C global warming is expected later this century^5,6,9^. COP26 also highlighted the vital importance of addressing global (in)equities and social (in)justices in tackling climate change^10^. But what is the human cost of climate change and who bears it? Existing estimates tend to be expressed in monetary terms^2^; tend to recognise impacts on the rich more than those on the poor (because the rich have more money to lose); and tend to value those living now over those living in the future (because future damages are subject to economic discounting). From an equity standpoint, this is unethical^3^ – when life or health are at stake, all people should be considered equal, whether rich or poor, alive or yet to be born.

A growing body of work considers how climate variability and climate change affect morbidity^11^ or mortality^12-15^. Here we take a complementary, ecological approach, considering exposure to less favourable climate conditions, defined as deviations of human population density with respect to climate from the historically-highly conserved distribution – the ‘human climate niche’^4^. The climate niche of species can be set by physiology^16^ and ecology^17^. For humans, the niche^4^ shows a primary peak of population density at MAT ∼13 °C and a secondary peak at ∼27 °C. The density of domesticated crops and livestock follow similar distributions^4^, as does GDP which shares the same independently identified^4,18^ primary temperature peak. Exposure outside the niche could result in increased morbidity, mortality, adaptation in place, or displacement (migration elsewhere). Humans have adapted physiologically and culturally to a wide range of local climates but despite this, mortality increases at both high and low temperatures^14,15^, consistent with the existence of a niche. Furthermore, high temperatures have been linked to decreased labour productivity^19^, decreased cognitive performance^20^, impaired learning^21^, adverse pregnancy outcomes^22^, decreased crop yield potential^11^, increased conflict^23-25^, migration^26^, and spread of infectious diseases^11,27,28^. Climate-related sources of harm not captured by the niche concept include sea level rise^29,30^.

## Results

### Reassessing the niche

First we re-examine how relative population density varies with mean annual temperature (MAT). Our previous work^4^ considered the 2015 population distribution under the 1960-1990 mean climate as a baseline (Extended Data Figure 1). Here we use the 1980 population distribution (total 4.4 billion) under the 1960-1990 mean climate (Figure 1a; ‘1980’) as the reference state. This is a more internally consistent approach, particularly as recent population growth biases towards hotter places. Applying a double-Gaussian fitting, the primary temperature peak is now larger and at slightly lower MAT (∼12 °C), in better agreement with reconstructions from 300 BP, 500 BP and 6000 BP (Extended Data Figure 1). The 1960-1990 interval was globally ∼0.3 °C warmer than the 1850-1900 ‘pre-industrial’ level, but closer to mean Holocene temperatures that supported civilisations as we know them (because 1850-1900 was at the end of the Little Ice Age). The smoothed double-Gaussian function fit (Figure 1a; ‘1980 fitted’) is referred to from hereon as the ‘temperature niche’. An updated ‘temperature-precipitation niche’ (additionally considering mean annual precipitation) was also calculated and considered in sensitivity analyses. By either definition, the niche is largely that of people dependent on farming. The niche of hunter-gatherers is likely broader^31-34^, as it is not constrained by the niches of domesticated species. This hypothesis is supported by the broader distribution of population density with respect to MAT reconstructed^4^ from the ArchaeoGlobe dataset for 6000 BP (when a smaller fraction of total population depended on farming) (Extended Data Figure 1b).

**Figure 1.**
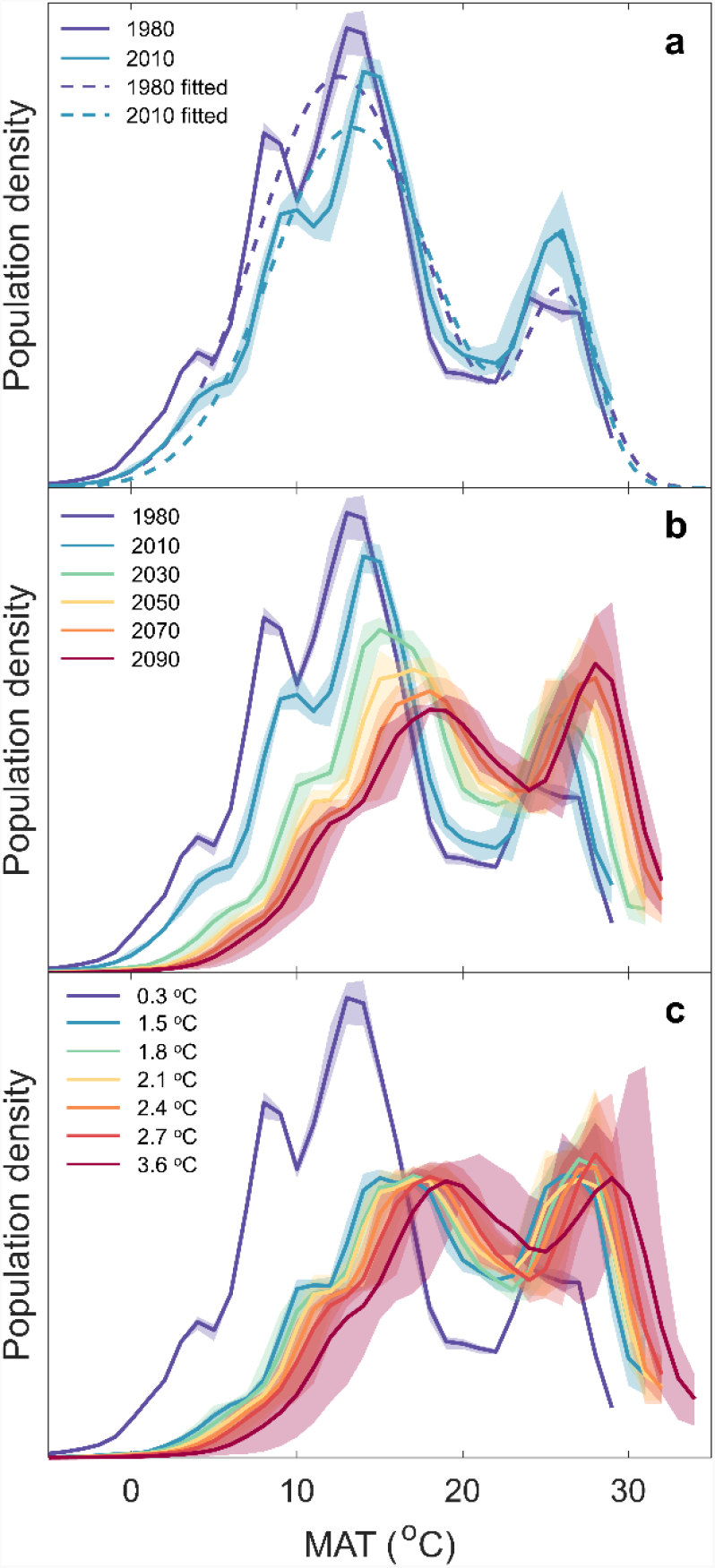
Changes in relative human population density with respect to Mean Annual Temperature (MAT). **a**. Observed changes from the reference distribution for 1980 population (4.4 billion) under 1960-1990 climate (0.3 °C global warming), to the 2010 population (6.9 billion) under 2000-2020 climate (1.0 °C global warming), together with smooth fitted functions (‘1980 fitted’ is defined as the temperature niche). **b**. Observed and projected future changes in population density with respect to MAT following SSP2-4.5 leading to ∼2.7 °C global warming and peak population 9.5 billion (see Extended Data Table 2 for global warming and population levels at each time). **c**. Projected population density with respect to MAT for a future world of 9.5 billion people under different levels of global warming (1.5, 1.8, 2.1, 2.4, 2.7, 3.6 °C), contrasted with the reference distribution (0.3 °C, 1980 population). Shaded regions correspond to 5-95% confidence intervals of the estimates.

### Calculating exposure

We provide three calculations of exposure outside of the temperature niche, due to: (i) unprecedented hot MAT, (ii) climate change only, or (iii) climate and demographic change (see Methods, Extended Data Figure 2). (i) The simplest approach^4^ just considers ‘hot exposure’ – i.e., how many people fall outside the hot edge of the temperature niche. This is calculated^4^ as the percentage of population exposed to MAT ≥29 °C. Only 0.3% of the 1980 population (12 million) experienced such conditions in the 1960-1990 climate. (ii) Exposure due to climate change alone^4^, considers all places where MAT changes to a value supporting lower relative population density according to the temperature niche: To calculate this^4^, we create a spatial ‘ideal distribution’ that maintains the historical distribution of relative population density with respect to MAT under a changed climate, and contrast this with the ‘reference distribution’ of population density with respect to the 1960-1990 climate. The difference between the ideal and reference distributions integrated across space gives the % of population exposed outside of the niche due to climate-only. (iii) Demographic change can also expose an increased density of population to a less favourable climate. To provide an upper estimate of population exposure (in %) due to both climate and demographic change we calculate the geographical distribution of projected population with respect to projected climate and contrast this ‘projected distribution’ with the ‘ideal distribution’.

### Changes up to present

We find that noticeable changes in the distribution of population density with respect to temperature have occurred due to climate and demographic changes from 1980 to 2010 (Figure 1a). Considering the 2010 population distribution (total 6.9 billion) under the observed 2000-2020 climate, global warming of 1.0 °C (0.7 °C above 1960-90) has shifted the primary peak of population density to slightly higher MAT (∼13 °C) compared to 1980, and the bias of population growth towards hot places has increased population density at the secondary (∼27 °C) peak. Greater observed global warming in the cooler higher northern latitudes than the tropics is visible in the changes to the distribution. Hot exposure (MAT ≥29 °C) doubled in percentage terms to 0.6% (41 million people), ∼8% of the global population have been left outside the historical niche due to climate change alone, and ∼12% from climate change plus demographic change. Thus, global warming of 0.7 °C since 1960-1990 has put ∼550 million people in less favourable climate conditions, with demographic change adding another ∼275 million.

### Future exposure

To estimate future exposure, we use an ensemble of eight climate model outputs (Extended Data Table 1) and corresponding population projections from four Shared Socio-economic Pathways^35^ (SSPs, Extended Data Table 2). SSP2-4.5 (‘middle of the road’) provides a useful reference scenario because it produces end-of-century (2081-2100) global warming 2.7 (2.1-3.5) °C corresponding to the 2.7 (2.0-3.6) °C expected under current policies^5^, and it captures largely unavoidable^36^ population growth towards a peak of ∼9.5 billion in 2070 (then declining to ∼9.0 billion in 2100). Climate change and population growth combine to shift relative population density to higher MAT (Figure 1b). Hot exposure (Figure 2a,d) becomes significant by 2030 at ∼4% or ∼0.3 billion as global warming reaches 1.5 °C, and it increases near linearly to ∼23% or ∼2.1 billion in 2090 under 2.7 °C global warming. People left outside the niche due to climate change alone (Figure 2b,e) reaches ∼14% or ∼1.2 billion by 2030, more than doubling to ∼29% or ∼2.7 billion in 2090. People left outside the niche from climate plus demographic change (Figure 2c,f) reaches ∼25% or ∼2.0 billion by 2030, and ∼40% or ∼3.7 billion by 2090.

**Figure 2.**
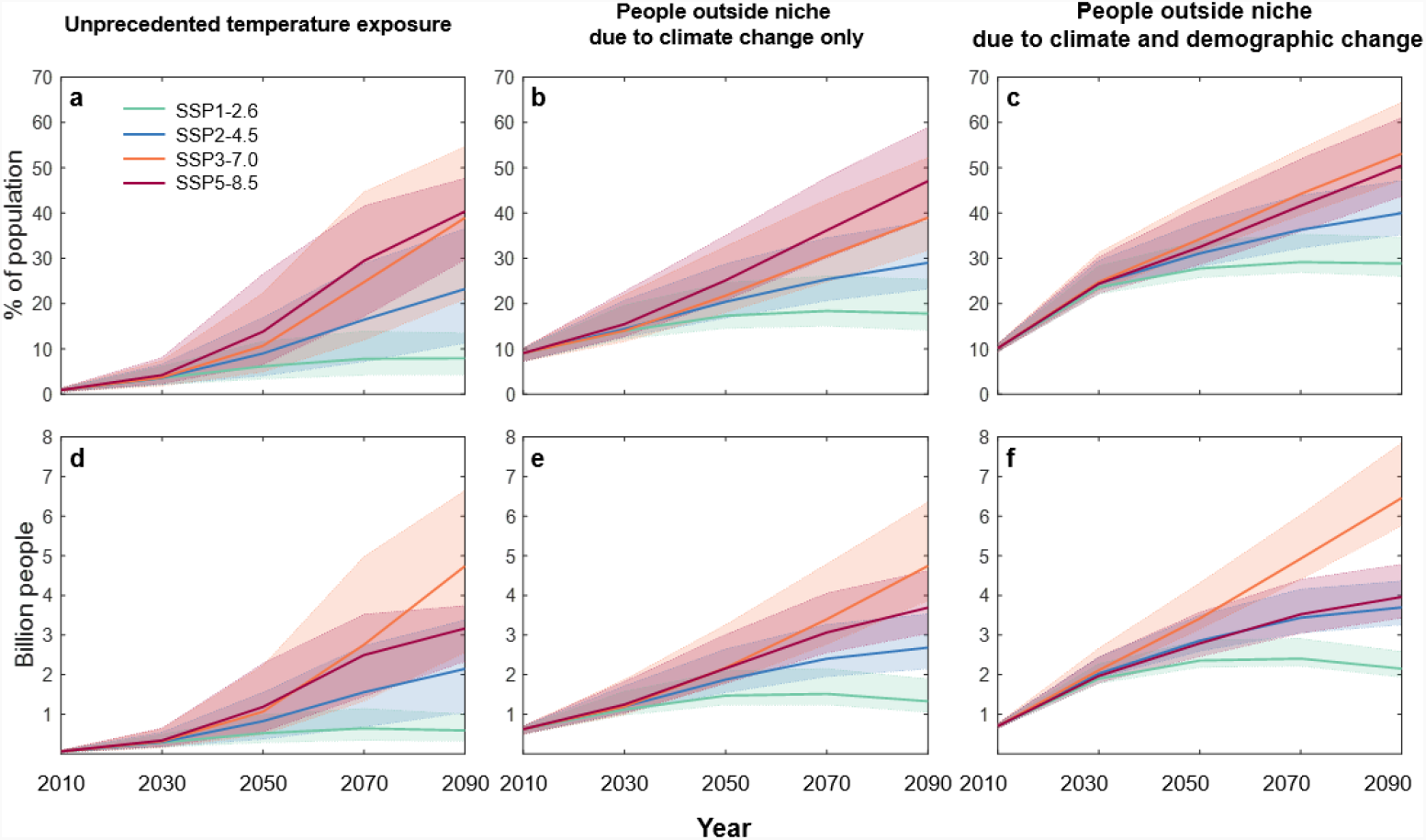
Population exposed outside of the temperature niche, following different Shared Socio-economic Pathways (SSPs): Top row (**a-c**): fraction of population (%); Bottom row (**d-f**): absolute population (billion people). **a, d**. Exposure to unprecedented hot mean annual temperature (MAT) ≥29 °C. **b, e**. People left outside of the niche due to climate change only. **c, f**. People left outside of the niche due to climate and demographic change. Calculations based on mean annual temperature (MAT) averaged over the 20-year intervals and population density distribution at the centre year of the corresponding intervals. The shaded regions correspond to 5-95% confidence intervals of the estimates.

### Variation across the SSPs

The other three Shared Socio-economic Pathways (SSPs) produce a wide range of global warming (2081-2100) from ∼1.8 (1.3-2.4) °C (SSP1-2.6) to ∼4.4 (3.3-5.7) °C (SSP5-8.5) and span a wide range of human development trajectories from population peaking at ∼8.5 billion then declining to ∼6.9 billion in 2100 (SSP1) to ongoing growth to ∼12.6 billion in 2100 (SSP3) (Extended Data Table 2). Both climate and demographic change alter the distribution of relative population density with respect to MAT (Extended Data Figure 3). By 2090, hot exposure reaches 8-40% or 0.6-4.7 billion across SSPs (Figure 2a,d). People left outside of the niche due to climate-only reaches 18-47% or 1.3-4.7 billion (Figure 2b,e). Adding in demographic change increases this to 29-53% or 2.2-6.5 billion (Figure 2c,f). Estimates of exposure outside the combined temperature-precipitation niche are roughly 20% greater than for the temperature niche alone (Extended Data Figure 4). SSP5-8.5 exposes the greatest proportion of population to unprecedented heat or being pushed out of the niche due to climate change alone, but SSP3-7.0 exposes the greatest proportion of population due to climate and demographic change combined, and the greatest absolute numbers across all three measures of exposure (Figure 2).

### Controlling for demography

Larger global populations following the SSPs place a greater proportion of people in hotter places, tending to leave more outside the historical niche (irrespective of climate change). To isolate the effects of climate policy and associated climate change on exposure, we fix the population and its distribution, exploring three different options: (i) 6.9 billion as in 2010, (i) 9.5 billion as in SSP2 in 2070, and (iii) 11.1 billion as in SSP3 in 2070. Having controlled for demography, global warming shifts the whole distribution of population density to higher temperatures (Figure 1c, Extended Data Figure 5). This results in linear relationships (Figure 3) between global warming and % of population exposed to unprecedented heat or left outside the niche from climate-only or climate plus demographic change. Hot exposure (Figure 3a) starts to become significant above the present level of ∼1.2 °C global warming and increases steeply at ∼12 % °C^-1^ (6.9 billion) to ∼17.5 % °C^-1^ (11.1 billion). People left outside the niche from climate change alone increases ∼12 % °C^-1^, above the baseline defined at 0.3 °C global warming (1960-1990) (Figure 3b). Factoring in demography, for a greater fixed population, the percent exposed is always greater, but the dependence on climate weakens somewhat towards ∼9 % °C^-1^ (for 11.1 billion). The relationships between global warming and exposure are all steeper for the temperature-precipitation niche (Extended Data Figure 6a). The mean temperature experienced by an average person increases with global warming in a manner invariant to demography at +1.5 °C °C^-1^ (Extended Data Figure 6b), consistent with observations and models that the land warms ∼1.5 times faster than the global average^37^.

**Figure 3.**
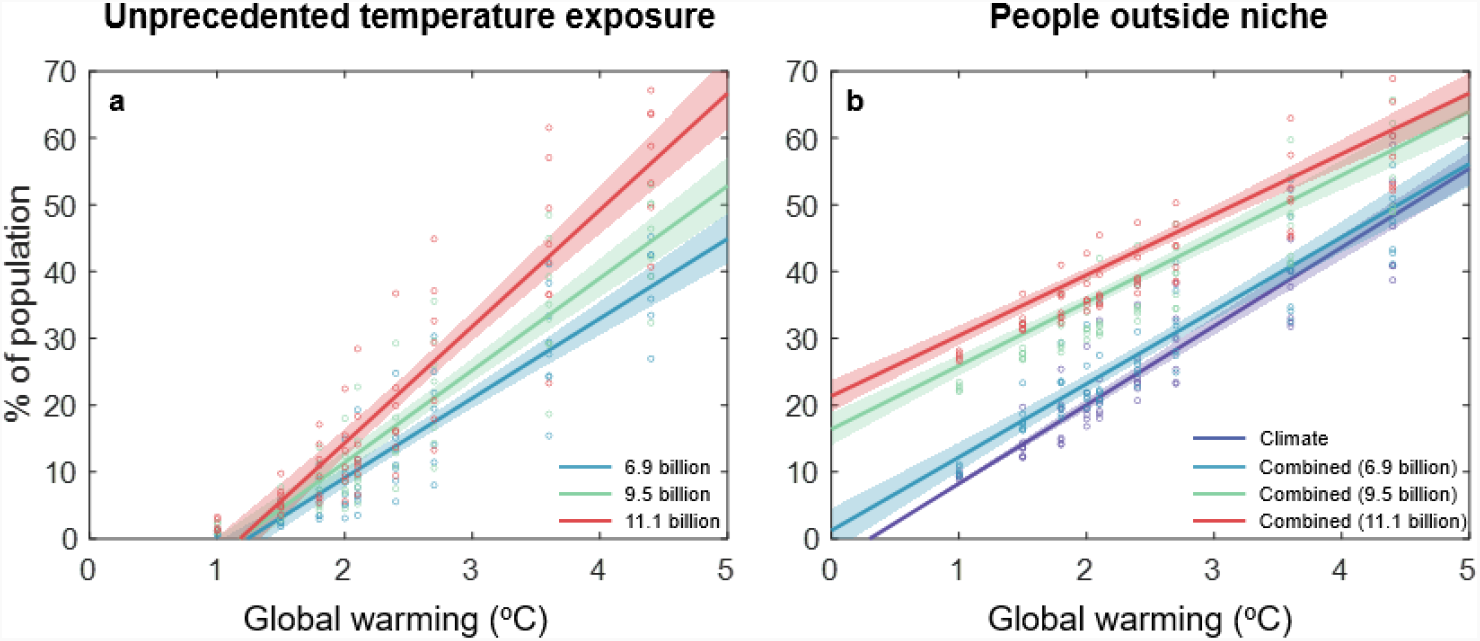
Relationships between global warming and population exposed outside of the temperature niche for different fixed population distributions: **a**. Population (%) exposed to mean annual temperature (MAT) ≥29 °C for the different population distributions: 6.9 billion (*n*=65, coefficient=11.9 % °C^-1^, *r*^2^=0.83); 9.5 billion (*n*=65, coefficient=13.8 % °C^-1^, *r*^2^=0.83); 11.1 billion (*n*=65, coefficient=17.5 % °C^-1^, *r*^2^=0.83). **b**. Population (%) put outside of the temperature niche due to climate change only (purple; *n*=65, coefficient=11.8 % °C^-1^; forcing intercept at 1960-1990 global warming of 0.3 °C), and due to the combined effects of climate and demographic change, for different fixed population distributions: 6.9 billion in 2010 (blue; *n*=65, coefficient=11.0 % °C^-1^, *r*^2^=0.83); 9.5 billion following SSP2 in 2070 (green; *n*=65, coefficient=9.5 % °C^-1^, *r*^2^=0.84), 11.1 billion following SSP3 in 2070 (red; *n*=65, coefficient=9.1 % °C^-1^, *r*^2^=0.84).

### Worst case scenarios

We now focus on a future world of 9.5 billion. When assessing risk it is important to consider worst case scenarios^38^. If the transient climate response to cumulative emissions is high, current policies could, in the worst case, lead to ∼3.6 °C end-of-century global warming^5^ (as projected under SSP3-7.0; Extended Data Table 2). This results in 33% (3.1 billion) hot exposed, 39% (3.7 billion) left outside the niche from climate-only, and 50% (4.8 billion) from climate plus demographic change (Figure 3). There also remains the possibility that climate policies are not enacted, and the world reverts to fossil-fuelled development (SSP5-8.5), leading to ∼4.4 °C end-of-century global warming. This gives 44% (4.2 billion) hot exposed, 49% (4.7 billion) left outside the niche from climate-only, and 59% (5.3 billion) from climate plus demography (Figure 3).

### Gains from strengthening climate policy

Having controlled for demography, strengthening climate policy reduces exposure (Figure 1c, 3), including to MAT ≥29 °C (Figure 4a,c), through reducing geographical movement of the temperature (Figure 4b,d) and temperature-precipitation (Extended Data Figure 7) niches. Following Climate Action Tracker’s projections^5^, different levels of policy ambition result in ∼0.3 °C changes in end-of-century global warming as follows: Current policies lead to ∼2.7 (2.0-3.6) °C; meeting current 2030 Nationally Determined Contributions (NDCs) (without long-term pledges) leads to ∼2.4 (1.9-3.0) °C; additional full implementation of submitted and binding long-term targets leads to ∼2.1 (1.7-2.6) °C; fully implementing all targets announced at COP26 leads to ∼1.8 (1.5-2.4) °C. Overall, going from ∼2.7 °C global warming under current policies to meeting the Paris Agreement 1.5 °C target reduces hot exposure from 21% to 5% (2.0 to 0.5 billion) (Figure 3a). It reduces population left outside the niche due to climate-only from 28% to 14% (2.7 to 1.3 billion), and it reduces population left outside the niche by climate plus demographic change from 42% to 31% (4.0 to 2.9 billion) (Figure 3b). Thus, each 0.3 °C decline in end-of-century warming reduces hot exposure by ∼4% or ∼380 million people, it reduces population left outside the niche due to climate-only by ∼3.5% or ∼330 million people, and population left outside the niche due to climate and demographic change by ∼3% or ∼260 million people.

**Figure 4.**
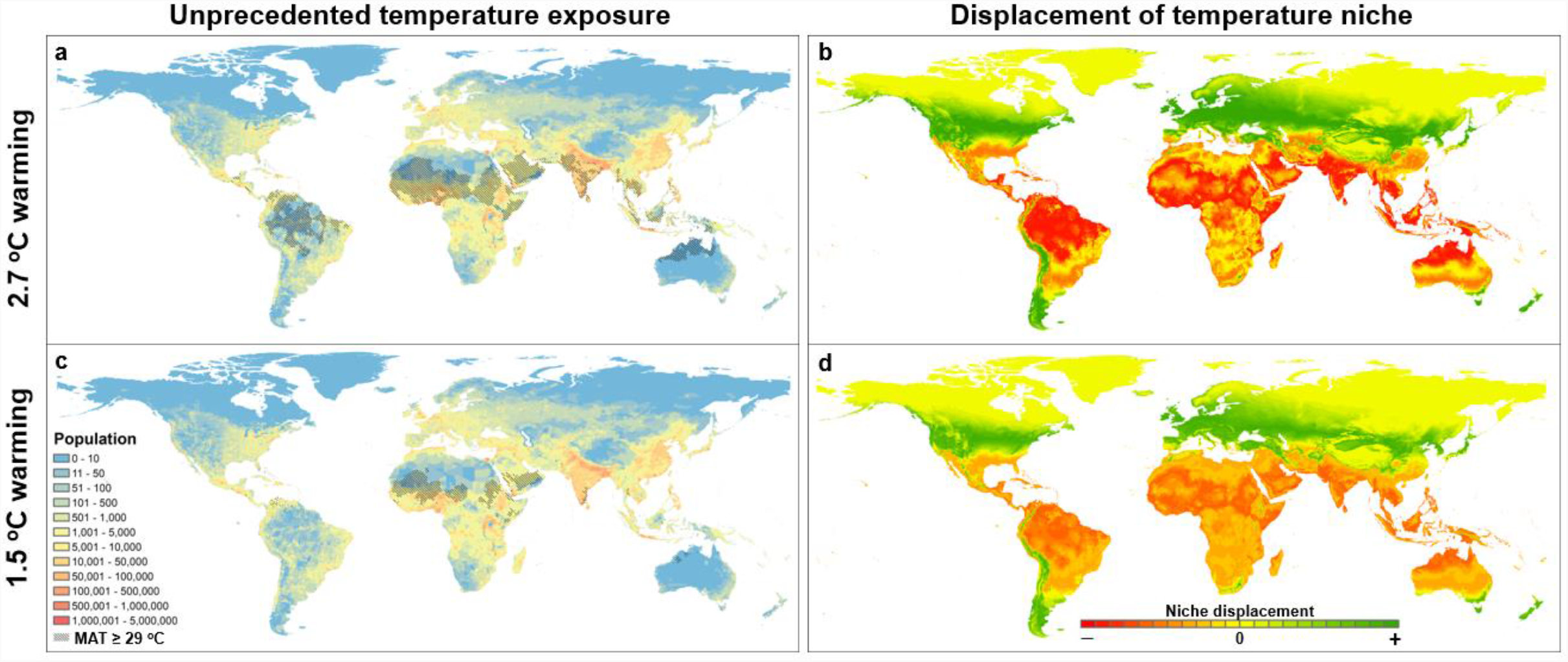
Global patterns of temperature niche change under **a**,**b**. 2.7°C and **c**,**d**. 1.5°C global warming. **a**,**c**. Regions exposed to unprecedented MAT ≥29 °C overlaid on population density (number in a ∼100 km^2^ grid cell) for a world of 9.5 billion (SSP2, 2070) under: **a**. 2.7°C global warming, **c**. 1.5°C global warming. **b**,**d**. Displacement of the temperature niche (red indicates a decrease in suitability, green an increase) under: **b**. 2.7°C global warming, **d**. 1.5°C global warming.

### Country-level exposure

We focus on hot exposure as the simplest and most conservative metric. The population exposed to unprecedented heat (MAT ≥29 °C) declines significantly for the most affected countries if global warming is reduced from ∼2.7 °C under current policies to meeting the 1.5 °C target (Figure 5). Assuming a future world of 9.5 billion, India has the greatest population exposed under 2.7 °C global warming, >600 million, but this reduces >6-fold to ∼90 million at 1.5 °C global warming. Nigeria has the second largest population exposed, >300 million under 2.7 °C global warming, but this reduces >7-fold to <40 million at 1.5 °C global warming. For third-ranked Indonesia, hot exposure reduces >20-fold from ∼100 million under 2.7 °C global warming, to <5 million at 1.5 °C global warming. For fourth and fifth-ranked Philippines and Pakistan with >80 million exposed under 2.7 °C global warming, there are even larger proportional reductions at 1.5 °C global warming. Sahelian-Saharan countries including Sudan (6th ranked) and Niger (7th) have a circa 2-fold reduction in exposure, because they still have a large fraction of land area exposed at 1.5 °C global warming (Extended Data Figure 8a). The fraction of land area exposed approaches 100% for several countries under 2.7 °C global warming (Extended Data Figure 8a). Brazil has the greatest absolute land area exposed under 2.7 °C global warming, despite almost no area being exposed at 1.5 °C, and Australia and India also experience massive increases in absolute area exposed (Figure 4a,b; Extended Data Figure 8b). (If the future population reaches 11.1 billion, the ranking of countries by population exposed remains similar, although the numbers exposed increases.) Those most exposed under 2.7 °C global warming come from nations that today are above the median poverty rate and below the median per capita emissions (Figure 6).

**Figure 5:**
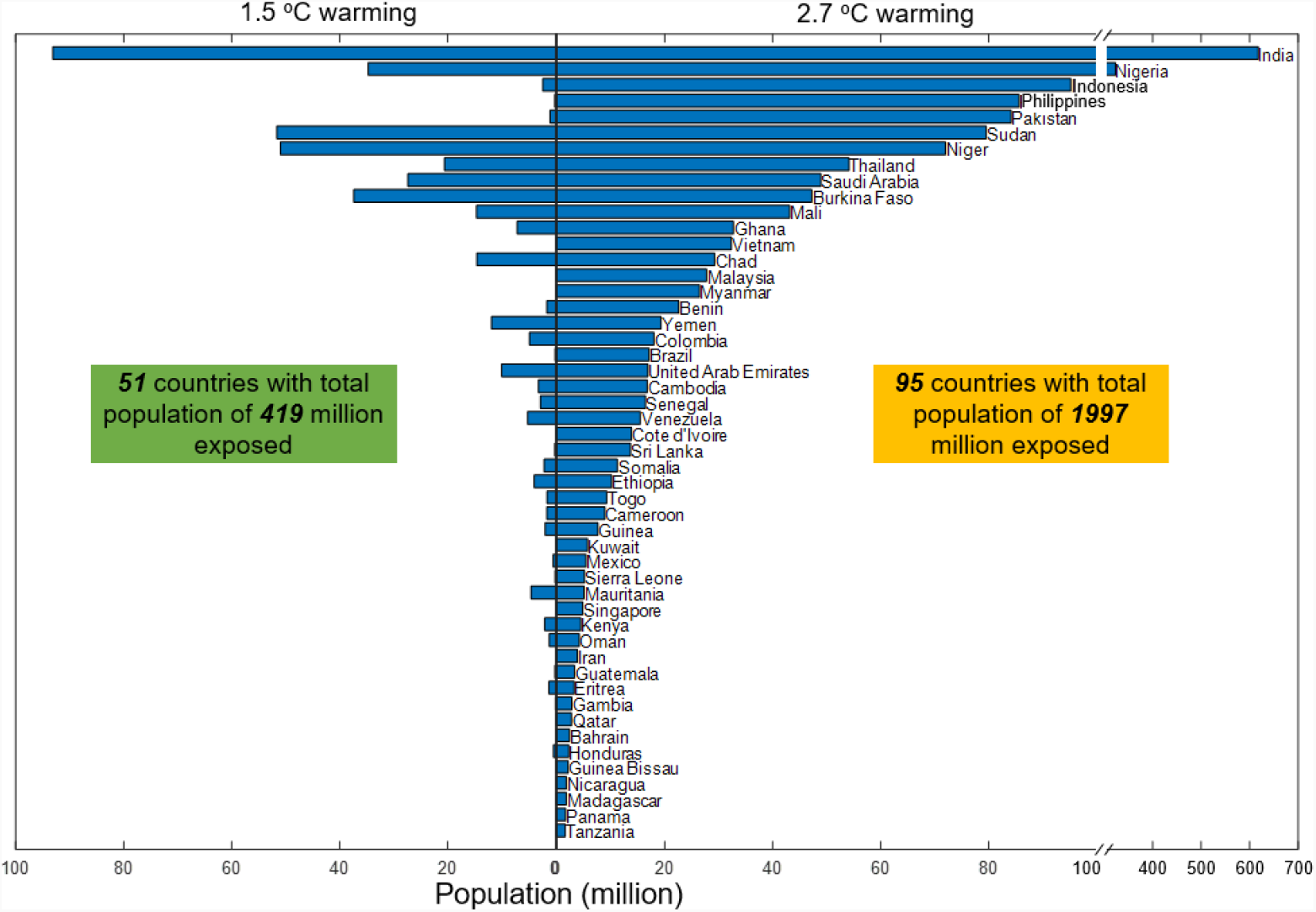
Country-level population exposed to unprecedented heat (MAT ≥29 °C) at 1.5 °C and 2.7 °C global warming. Showing the top 50 countries (ranked under 2.7 °C global warming) in a world of 9.5 billion people (around 2070 under SSP2). Note the break in the x-axis for the top 2 countries.

**Figure 6.**
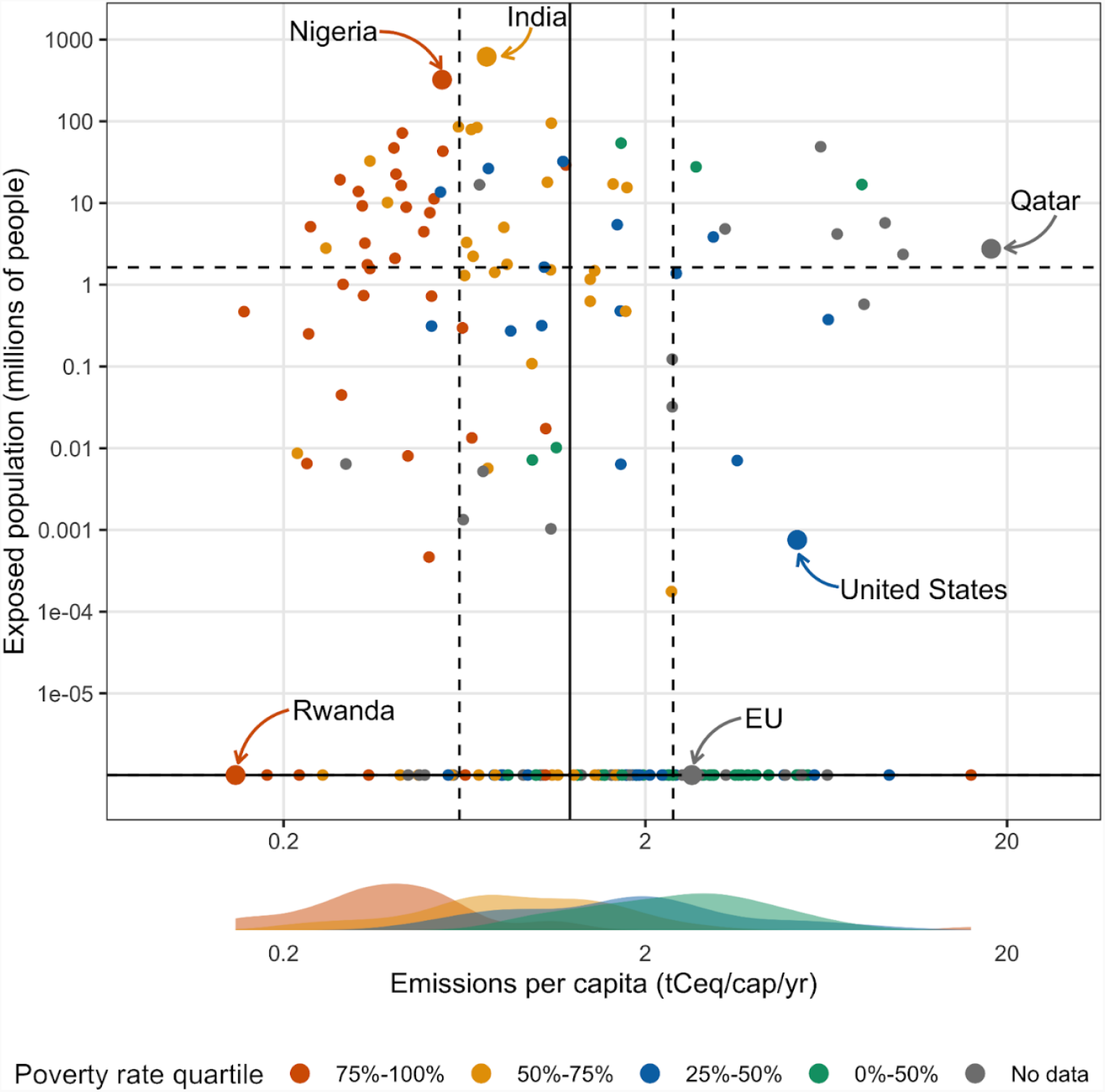
Country-level per capita greenhouse gas emissions^40^ related to population exposed to unprecedented heat (MAT ≥29 °C) at 2.7 °C global warming (Figure 5) and poverty rate^50^. Solid lines show the median (50% quantile) and dashed lines the 25% and 75% quantiles for emissions and heat exposure. Points are coloured by quartile of the poverty rate distribution, where poverty rate is defined as % of national population below the $1.90 poverty line. The density plots at the bottom show the distribution of emissions per capita for each poverty rate quartile.

### Relating present emissions to future exposure

Above the present level of 1.2 °C global warming, the increase in hot exposure of 13.8 % °C^-1^ for a future world of ∼9.5 billion people, corresponds to 1.31×10^9^ cap °C^-1^. The established relationship^39^ of cumulative emissions to transient global warming is ∼1.65 (1.0-2.3) °C EgC^-1^. Therefore 1 person will be exposed to unprecedented heat (MAT ≥29 °C) for every ∼460 (330-760) tC emitted. Present (2018 data) global mean per capita CO_2_-equivalent emissions^40^ (production-based) are 1.8 tC_eq_ cap^-1^ yr^-1^. Thus, during their lifetimes (72.6 years) ∼3.5 global average citizens today (less than the average household of 4.9 people) emit enough carbon to expose one future person to unprecedented heat. Citizens in richer countries generally have higher emissions^40^, e.g. EU 2.4 tC_eq_ cap^-1^ yr^-1^, US 5.3 tC_eq_ cap^-1^ yr^-1^, Qatar 18 tC_eq_ cap^-1^ yr^-1^ (Figure 6) (and consumption-based emissions are even higher). Thus, ∼2.7 average EU citizens or ∼1.2 average US citizens emit enough carbon in their lifetimes to expose 1 future person to unprecedented heat, and the average citizen of Qatar emits enough carbon in their lifetime to expose ∼2.8 future people to unprecedented heat. Those future people tend to be in nations that today have per capita emissions around the 25% quantile (Figure 6), including the two countries with the greatest population exposed: India 0.73 tC_eq_ cap^-1^ yr^-1^ and Nigeria 0.55 tC_eq_ cap^-1^ yr^-1^. We estimate that the average future person exposed to unprecedented heat comes from a place where today per capita emissions are approximately half (56%) of the global average (or 52% in a world of 11.1 billion people).

## Discussion

Our estimate that global warming since 1960-1990 has put more than half a billion people outside the historical climate niche is consistent with attributable impacts of climate change affecting 85% of the world’s population^41^. Above the present level of ∼1.2 °C global warming, predicted exposure to unprecedented MAT ≥29 °C increases markedly. This is consistent with extreme humid heat having more than doubled in frequency^42^ since 1979, with exposure in urban areas increasing for 23% of the world’s population^43^ from 1983 to 2016 (due also to growing urban heat islands), and the total urban population exposed tripling^43^ (due also to demographic change). Extreme humid heat can be fatal^44^ for more vulnerable individuals^45^, and is associated with labour loss of 148 million full time equivalent jobs^19^. Further work should examine the relationship between unprecedented MAT ≥29 °C and exposure to extreme heat. Both India and Nigeria already show ‘hotspots’ of increased exposure to extreme heat due predominantly to warming^43^, consistent with our prediction that they are at greatest future risk (Figure 5). These and other emerging economies (e.g., Indonesia, Pakistan, Thailand) dominate the total population exposed to unprecedented heat in a 2.7 °C warmer world (Figure 5). Their climate policy commitments also play a significant role in determining end-of-century global warming^9^.

The huge numbers of humans left out of the climate niche in our future projections warrant critical evaluation. Combined effects of climate and demographic change are upper estimates. This is because at any given time the method limits absolute population density of the (currently secondary) higher-temperature peak based on absolute population density of the (currently primary) lower-temperature peak. Yet absolute population density is allowed to vary (everywhere) over time. (This is not an issue for the climate-only or hot-exposure estimates.) Nevertheless, a bias of population growth to hot places clearly increases the proportion (as well as the absolute number) of people exposed to harm from high temperatures^46^.

Overall, our results illustrate the huge potential human cost and the great inequity of climate change – without having considered exposure to e.g., sea-level rise^29,30^. They also inform discussions of loss and damage^47,48^. The worst-case scenarios of ∼3.6 °C or even ∼4.4 °C global warming could put half of the world population outside the historical climate niche, posing an existential risk. The ∼2.7 °C global warming expected under current policies puts around a third of world population outside the niche. It exposes almost the entire area of some countries (e.g., Burkina Faso, Mali) to unprecedented heat, including some Small Island Developing States (SIDS) (e.g., Aruba, Netherlands Antilles) (Extended Data Figure 8b) – a group with members already facing an existential risk from sea-level rise. The gains from fully implementing all policy targets announced at COP26 and limiting global warming to ∼1.8 °C are considerable but would still leave nearly 10% of people exposed to MAT ≥29 °C. Meeting the goal of the Paris Agreement to limit global warming to 1.5 °C, halves exposure outside of the historical niche relative to current policies, and limits those exposed to unprecedented hot temperatures to 5% of people. This still leaves several least-developed countries (e.g., Sudan, Niger, Burkina Faso, Mali) with large populations exposed (Figure 5), adding adaptation challenges to an existing climate investment trap^49^. Nevertheless, our results show the huge potential for more decisive climate policy to limit the human costs and inequities of climate change.

## Methods

### Reassessing the climate niche

We plot the running mean of population density against mean annual temperature (MAT), with a step of 1 °C and a bin size of 2 °C, and then apply double-Gaussian fitting to the resulting curve^4^. Our previous work^4^ assessed the human temperature niche by quantifying the 2015 population distribution in relation to the 1960-1990 MAT (Extended Data Figure 1; ‘Old reference’). Here we re-assessed the niche changing the data to the 1980 population distribution (total 4.4 billion) under the 1960-1990 MAT, for greater internal consistency (Figure 1a, Extended Data Figure 1; ‘1980’). This is important because there has been significant population growth between 1980 and 2015 with a distinct bias to hotter places. The 1980 population distribution data was obtained from the HYDE (History database of the Global Environment) 3.2 database^51^. The ensemble mean 1960-1990 climate and associated uncertainty (5th/95th percentiles) were calculated from three sources: (1) WorldClim v1.4 data^52^, (2) CRU TS v.4.05 monthly data^53,54^ (available at https://crudata.uea.ac.uk/cru/data/hrg/), (3) NASA GLDAS-2.1 (Global Land Data Assimilation System) 3-hourly data^55^ (available at https://developers.google.com/earth-engine/datasets/catalog/NASA_GLDAS_V021_NOAH_G025_T3H). A revised temperature-precipitation niche was also calculated from both MAT and mean annual precipitation (MAP), following the methods in ref. ^4^, but using the 1980 population distribution with the 1960-1990 mean climate. This is considered in sensitivity analyses.

### Projecting the niche

Hot exposure is calculated (as previously^4^) for a given climate and population distribution as the percentage of people exposed to mean annual temperature (MAT) ≥29 °C, from a direct spatial comparison of MAT and population distributions (without any smoothing). The MAT ≥29 °C threshold was chosen as only 0.3% of the 1980 population (12 million) experienced such conditions in the 1960-1990 climate. To separate the effects of climate and demographic changes on geographic displacement of the human climate niche, we consider (Extended Data Figure 2): (1) the geographic distribution of the reference niche (‘reference distribution’), (2) projecting the reference niche function to the geographic distribution of present/future climate (‘ideal distribution’), and (3) the geographically projected ‘assumed distribution’, of present/future population with respect to present/future climate conditions. Here (2) minus (1) gives the effect of climate change only (as previously^4^), and (3) minus (2) gives the combined effect of climate and demographic change.

### Changes up to present

To calculate changes up to (near) present we construct an ensemble mean 2000-2020 climate and associated uncertainty (5th/95th percentiles) from five sources: (1) CRU TS v.4.05 monthly data^53,54^; (2) NASA GLDAS-2.1 (Global Land Data Assimilation System) 3-hourly data^55^; (3) ECMWF ERA5-Land monthly averaged Climate Reanalysis data^56^ (available at https://developers.google.com/earth-engine/datasets/catalog/ECMWF_ERA5_LAND_MONTHLY); (4) NASA FLDAS (Famine Early Warning Systems Network Land Data Assimilation System) monthly data^57,58^ (available at https://developers.google.com/earth-engine/datasets/catalog/NASA_FLDAS_NOAH01_C_GL_M_V001); (5) NCEP CFSV2 (Climate Forecast System Version 2) 6-hourly data^59^ (available at https://developers.google.com/earth-engine/datasets/catalog/NOAA_CFSV2_FOR6H). Each climate dataset is aggregated to calculate mean annual temperature and precipitation. The 2000-2020 climate represents 1.0 °C global warming relative to the pre-industrial level. The 2010 population distribution data was obtained from the HYDE (History database of the Global Environment) 3.2 database^51^. We followed the methods described above to calculate exposure.

### Future projections

We used projected climate and population distribution under four different Shared Socio-economic Pathways (SSPs), which combine different demographic and emissions projections under consistent storylines: SSP1-2.6 (sustainability), SSP2-4.5 (middle of the road), SSP3-7.0 (regional rivalry), and SSP5-8.5 (fossil-fuelled development). We focused on 20-year mean climate states for 2020-2040, 2040-2060, 2060-2080, and 2080-2100, and the projected population distribution data of 2030, 2050, 2070 and 2090 to represent average demographic conditions of corresponding time periods. We obtained downscaled Coupled Model Intercomparison Project phase 6 (CMIP6) climate data available from WorldClim v2.0 at 0.0833 degree (∼10 km) resolution (available at https://worldclim.org), which restricts us to up to eight CMIP6 models (Extended Data Table 1). We obtained SSP population data at 1-km resolution from the spatial population scenarios dataset^60,61^ (available at https://www.cgd.ucar.edu/iam/modeling/spatial-population-scenarios.html). We aggregated the population data to a consistent resolution of 0.0833 degree (∼10 km) to match the climate data and our previous analyses. We combine results across climate models to create a multi-model ensemble mean, and a 5-95% confidence interval, recognising that the number of models available varies somewhat between SSPs and time-slices (Extended Data Table 1). To this end, we apply the MAT data of each climate model to plot population density against MAT and then combine the resulting curves to calculate the mean, and 5th and 95th percentiles.

To control for demography and thus isolate the effects of climate policy and associated climate change on exposure, we consider three different fixed populations and their spatial distributions: (i) 6.9 billion as in 2010, (i) 9.5 billion following SSP2 in 2070, and (iii) 11.1 billion following SSP3 in 2070. These are combined with the observed (2000-2020) 1.0 °C global warming and with different future levels of global warming (1.5, 1.8, 2.0, 2.1, 2.4, 2.7, 3.6, 4.4 °C) corresponding to different 20-year climate averages from different SSPs (Figure 1c, 3, Extended Data Figure 5). Global warming of 1.5 °C and 2.0 °C are considered because of their relevance to the Paris Agreement. 1.8, 2.1, 2.4 and 2.7 °C are chosen as best estimates of end-of-century global warming corresponding to different policy assumptions, taken from the ‘Climate Action Tracker’^5^, which uses an ensemble of runs of the MAGICC6 model that in turn emulates different general circulation models (GCMs) from CMIP6. Global warming of 3.6 and 4.4 °C are chosen as worst case scenarios that also enable examining the shape of relationships between global warming and population exposure. 20-year SSP intervals corresponding to these different levels of global warming are chosen based on mean global warming levels from the CMIP6 model ensemble given in Table SPM.1 of the Sixth Assessment Report (AR6) of the Intergovernmental Panel on Climate Change^62^ (IPCC). We try to match to warming in 2081-2100, but where earlier time intervals must be used this should have little effect on the results because the spatial pattern of temperature change is highly conserved on the century timescale. The different combinations are: 1.5 °C = SSP1-2.6 in 2021-2040; 1.8 °C = SSP1-2.6 in 2081-2100; 2.0 °C = SSP2-4.5 in 2041-2060; 2.1 °C = SSP3-7.0 in 2041-2060; 2.4 °C = SSP5-8.5 in 2041-2060; 2.7 °C = SSP2-4.5 in 2081-2100; 3.6 °C = SSP3-7.0 in 2081-2100; 4.4 °C = SSP5-8.5 in 2081-2100. For the same time interval and SSP, different CMIP6 models can give different levels of global warming due to differing climate sensitivity. This is apparent in the spread of population exposure results for individual models (open circles in Figure 3, Extended Data Figure 6). However, we checked that global warming in the multi-model ensemble mean of the CMIP6 models we consider (Extended Data Table 1) matches that of the larger CMIP6 ensemble (Table SPM.1 of IPCC AR6).

### Country-level estimates

Results for hot exposure were disaggregated to country scale for 2.7 °C and 1.5 °C global warming and populations of 9.5 or 11.1 billion, using GIS data for country boundaries from the World Borders Dataset (https://thematicmapping.org/downloads/world_borders.php).

### Emissions and poverty rate of those exposed

Using the country-level breakdown of exposure to unprecedented heat in a 2.7 °C warmer world with 9.5 billion people (Figure 5) we calculated a weighted average for number of people exposed multiplied by percentage of global average emissions per capita today. This uses production-based, country-level CO_2_-equivalent greenhouse gas emissions from the emissions database for global atmospheric research^40^, for which 2018 is the latest year. The calculation was also done for country-level exposure in a 2.7 °C warmer world of 11.1 billion. Consumption-based emissions (accounting for trade) tend to be lower than production-based emissions in poorer countries and higher in richer countries. This would increase the inequity already apparent in the results. We also examined poverty rate defined as the percentage of population per country below the $1.90 poverty line, using the interpolated data for 2019 from the World Bank’s Poverty and Inequality Platform^50^ (https://pip.worldbank.org/home). The resulting distribution is heavily skewed with 25% quantile = 0.26%, 50% quantile = 1.79%, 75% quantile = 20%.

## Data availability

All data analysed during this study are available at the URLs given (above), with the WorldClim v1.4 data available within https://doi.org/10.5061/dryad.fj6q573q7. All data generated during this study are available from corresponding author C.X. and will be deposited in dryad before final publication.

## Code availability

Code used for the analysis is available from corresponding author C.X. and will be deposited in dryad before final publication.

## Acknowledgements

We thank all the data providers. T.M.L., J.F.A. and A.G. are supported by the Open Society Foundations. T.M.L. is supported by a Turing Fellowship. C.X. is supported by the National Natural Science Foundation of China (32061143014) and the Fundamental Research Funds for the Central Universities. This work is part of the Earth Commission which is hosted by Future Earth and is the science component of the Global Commons Alliance. The Global Commons Alliance is a sponsored project of Rockefeller Philanthropy Advisors, with support from Oak Foundation, MAVA, Porticus, Gordon and Betty Moore Foundation, Herlin Foundation and the Global Environment Facility. J.C.S. is supported by VILLUM Investigator project “Biodiversity Dynamics in a Changing World” funded by VILLUM FONDEN (grant 16549). M.S. is supported by an ERC Advanced Grant and a Spinoza award.

## Author contributions

T.M.L., C.X. and M.S. designed the study. C.X. performed the climate niche analyses with input from T.M.L.; T.M.L. and J.F.A. related present emissions to future exposure. C.X. and J.F.A. produced the figures with input from T.M.L., S.L., B.S. and C.Z.; T.M.L. wrote the paper with input from all authors.

## Extended Data Figures

**Extended Data Figure 1.**
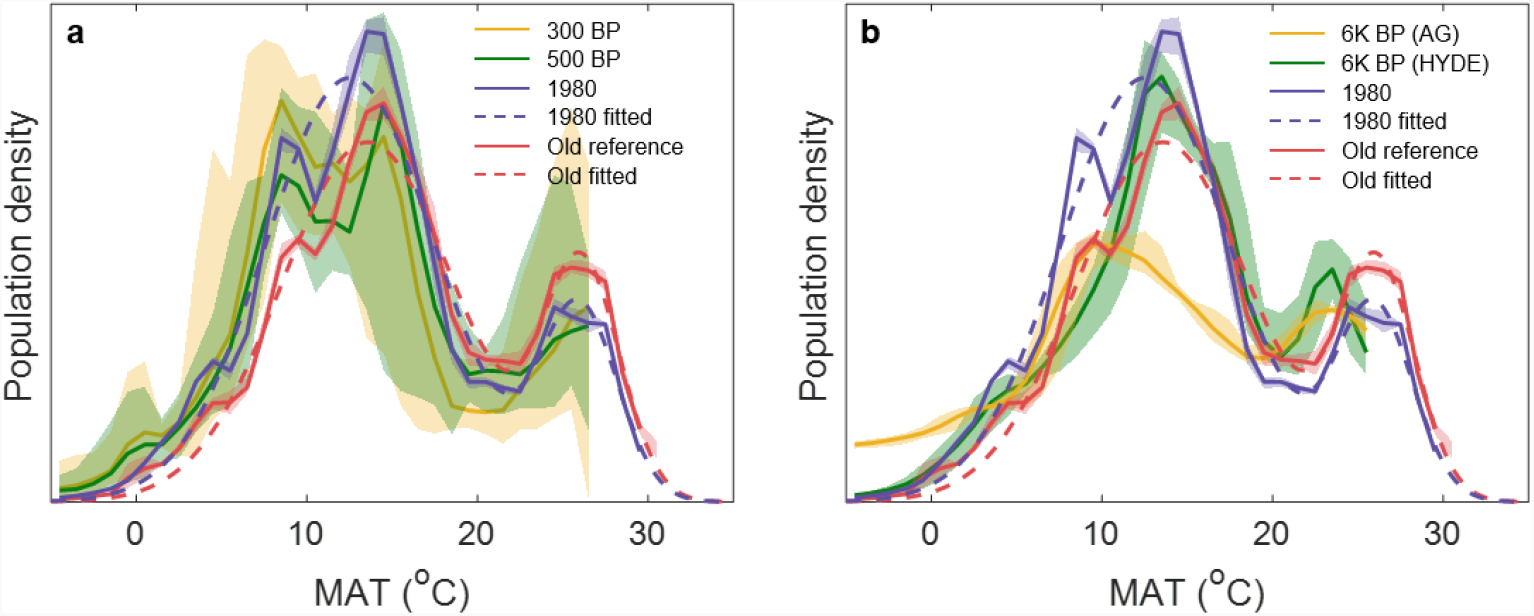
Relative human population density with respect to Mean Annual Temperature (MAT). Reconstructions for **a**. 300 BP, 500 BP (population data from HYDE database), and **b**. 6000 BP with population data from ArchaeoGlobe (AG) or HYDE, compared to the 1960-1990 climate (∼0.3 °C above pre-industrial) with 2015 population distribution (‘Old reference’) or 1980 population distribution (‘1980’; as in Figure 1a), and the smooth fitted functions for the temperature niche used previously (‘Old fitted’) and here (‘1980 fitted’; as in Figure 1a) for future projections. The shaded regions correspond to 5-95% confidence intervals of the estimates.

**Extended Data Figure 2.**
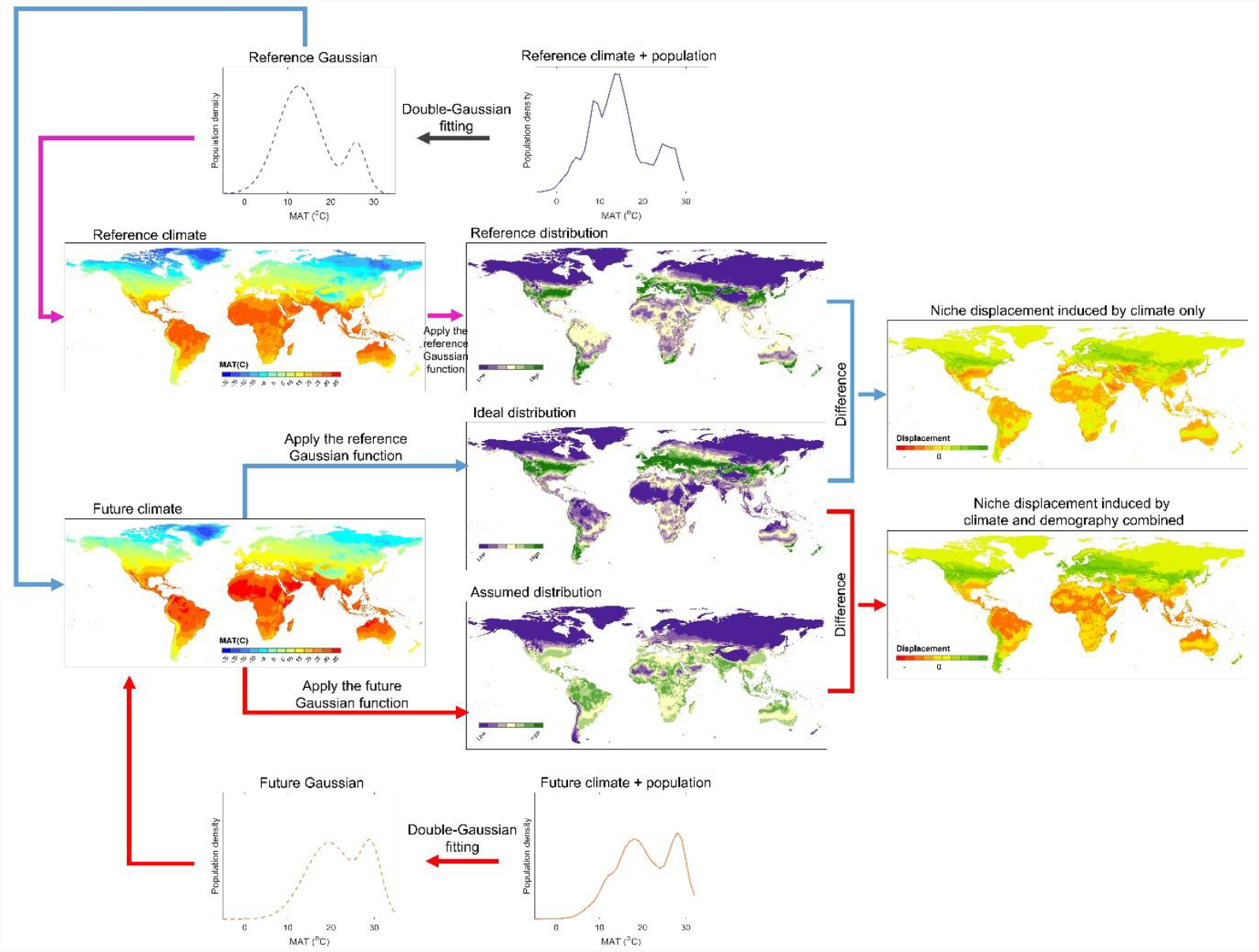
Illustration of workflow for quantifying displacement of the human climate niche due to climate change only or climate and demographic change – here for the temperature niche (but the same approach is used for the temperature-precipitation niche).

**Extended Data Figure 3.**
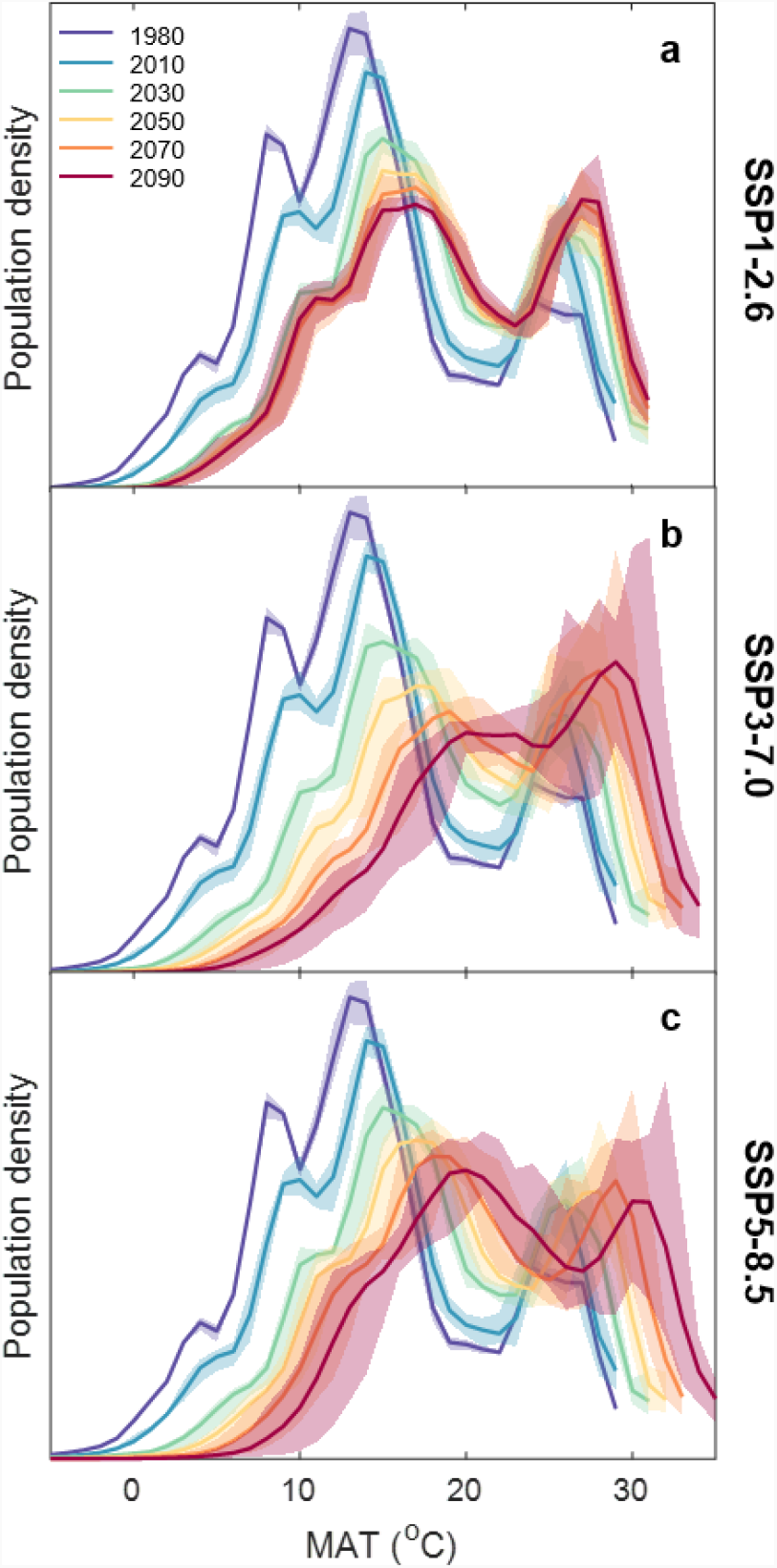
Observed and projected future changes in human population density with respect to Mean Annual Temperature (MAT), following different Shared Socio-economic Pathways (SSPs): **a**. SSP1-2.6 leading to ∼1.8 °C global warming with a peak of 8.5 billion people. **b**. SSP3-7.0 scenario leading to ∼3.6 °C global warming and 12.1 billion people. **c**. SSP5-8.5 scenario leading to ∼4.4 °C global warming and a peak of 8.6 billion people. (The SSP2-4.5 scenario is shown in Figure 1b.) For each SSP and 20-year averaged climate interval, global warming and corresponding population levels (for the central year) are summarised in Extended Data Table 2. Shaded regions correspond to 5-95% confidence intervals of the estimates.

**Extended Data Figure 4.**
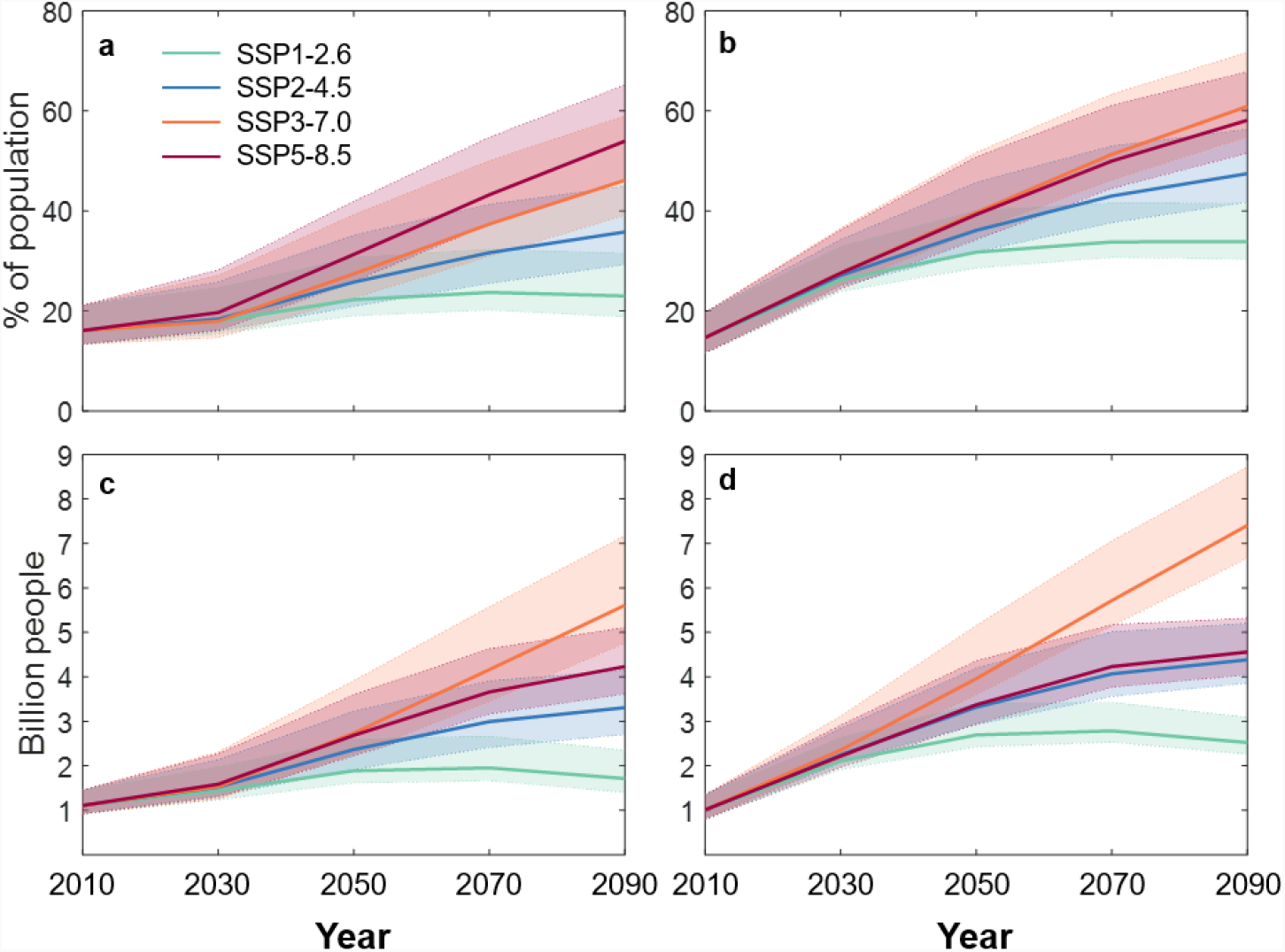
Population exposed outside of the temperature-precipitation niche, following different Shared Socio-economic Pathways (SSPs): **a**,**b**. Fraction of population (%) left outside of the niche due to: **a**. climate change only. **b**. climate and demographic change. **c**,**d**. Absolute number left outside of the niche due to: **c**. climate change only. **d**. climate and demographic change. Calculations based on mean annual temperature (MAT) averaged over the 20-year intervals and population density distribution at the centre year of the corresponding intervals. The shaded regions correspond to 5-95% confidence intervals of the estimates. (Note that the population exposed to unprecedented hot MAT ≥29 °C is unaltered by considering precipitation changes.)

**Extended Data Figure 5.**
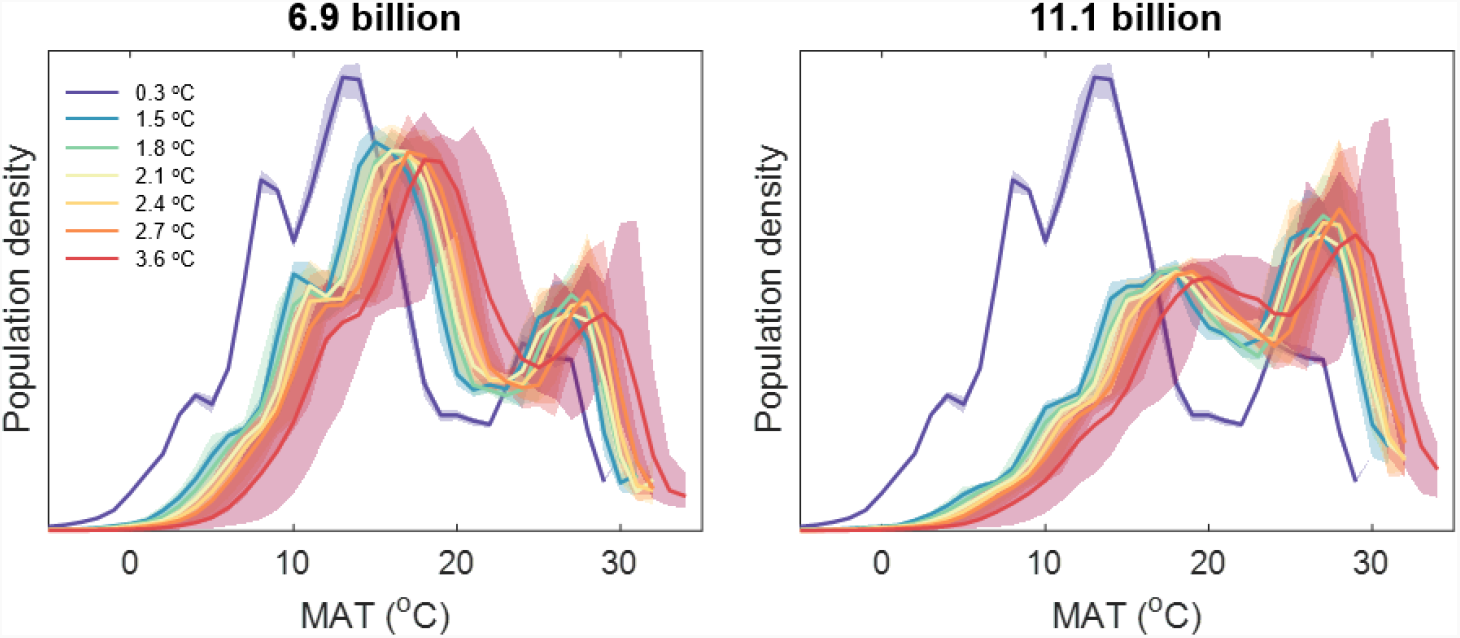
Changes in human population density with respect to Mean Annual Temperature (MAT) for different fixed population distributions and levels of global warming. The population distributions are: **a**. 6.9 billion in 2010, **b**. 11.1 billion under SSP3 in 2070 (9.5 billion under SSP2 in 2070 is shown in Figure 1c). See Methods for the combinations of SSP and 20-year time interval representing different global warming levels.

**Extended Data Figure 6.**
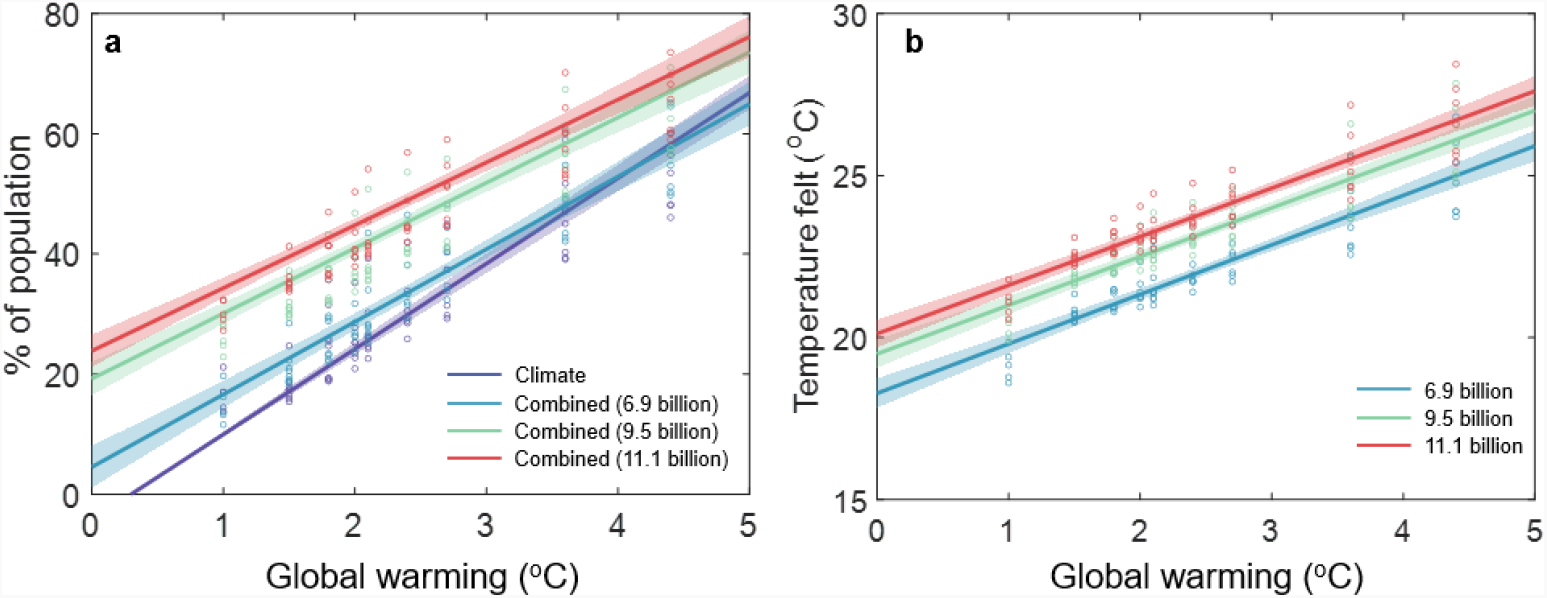
Near linear relationships between global warming and niche displacement and between global warming and average temperature experienced. **a**. Near linear relationship between global warming and temperature-precipitation niche displacement (%) due to climate-only and due to climate plus demographic change. Linear regression results: Climate-only (*n*=65, coefficient=14.2 % °C^-1^; forcing intercept at 1960-1990 global warming of 0.3 °C); Combined 6.9 billion (*n*=65, coefficient=12.0 % °C^-1^, *r*^2^=0.84); Combined 9.5 billion (*n*=65, coefficient=10.9 % °C^-1^, *r*^2^=0.84); Combined 11.1 billion (*n*=65, coefficient=10.5 % °C^-1^, *r*^2^=0.84). **b**. Mean annual temperature felt by an average person for different levels of global warming for fixed population distributions. Linear regression results: 6.9 billion (*n*=65, coefficient=1.53 °C °C^-1^, *r*^2^=0.83); 9.5 billion (*n*=65, coefficient=1.50 °C °C^-1^, *r*^2^=0.84); 11.1 billion (*n*=65, coefficient=1.50 °C °C^-1^, *r*^2^=0.84).

**Extended Data Figure 7.**
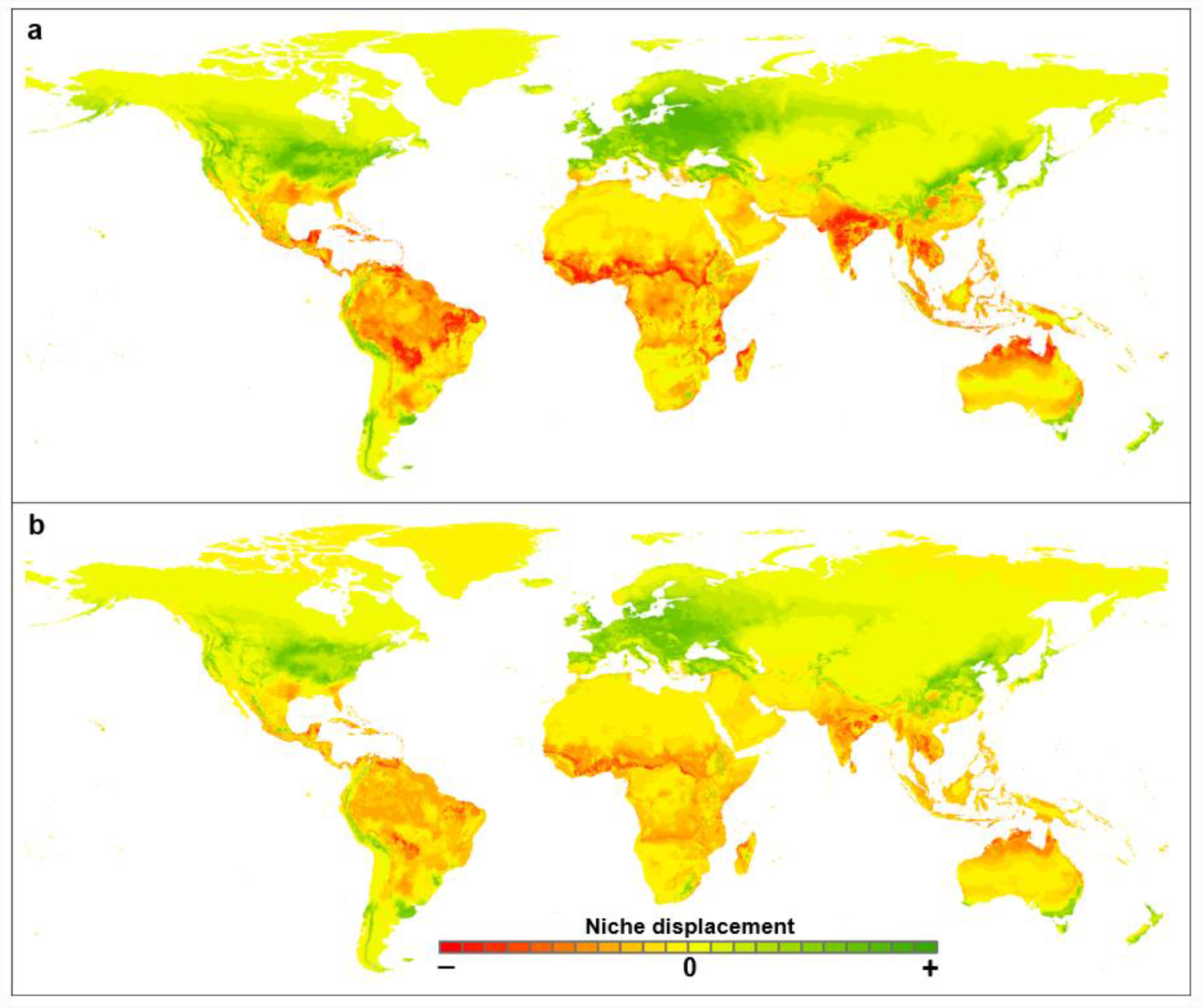
Displacement of the temperature-precipitation niche under **a**. 2.7°C global warming due to current policies, **b**. 1.5°C global warming meeting the Paris Agreement. Red indicates a decrease in suitability, green an increase.

**Extended Data Figure 8.**
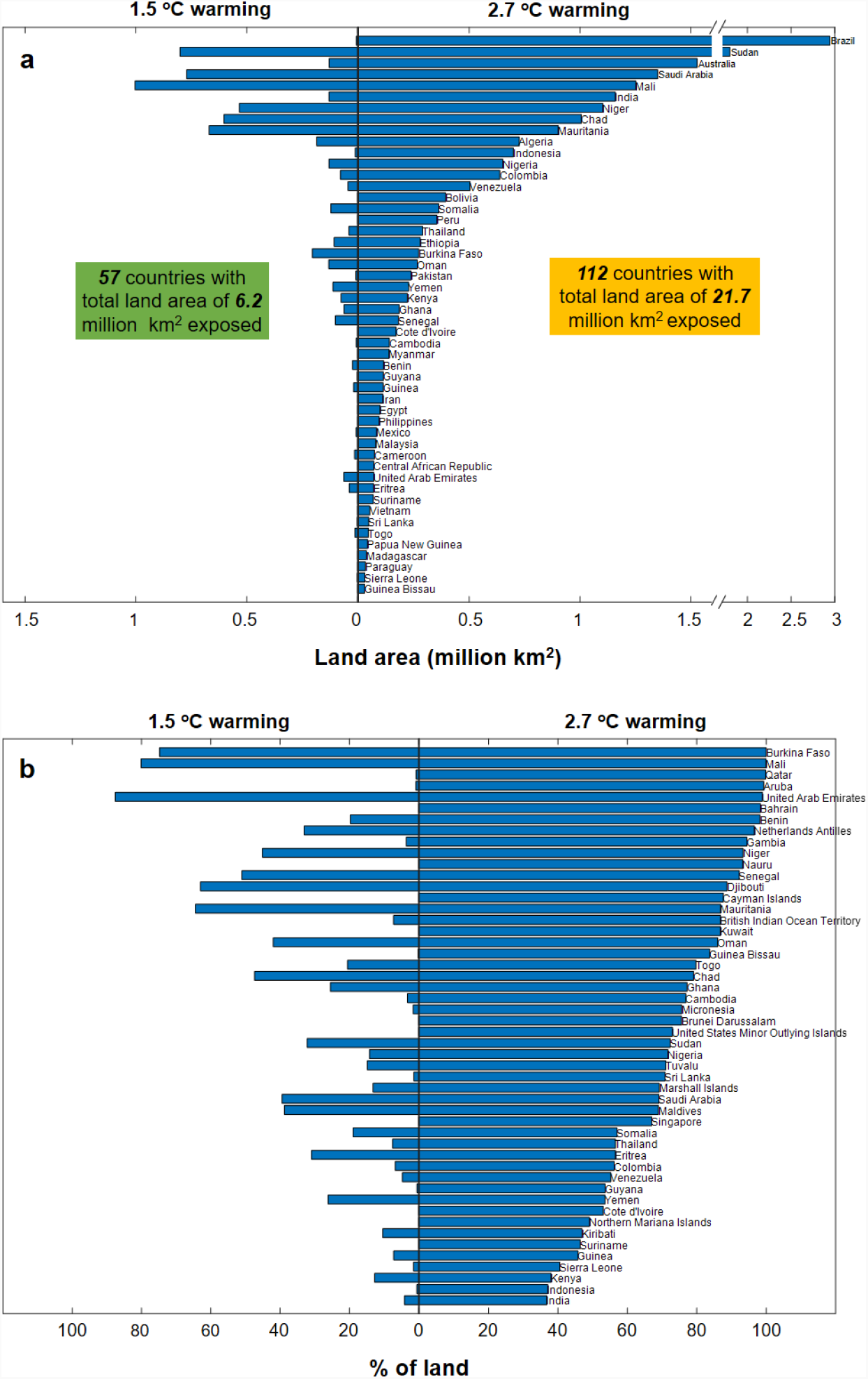
Land area exposed to unprecedented heat (MAT ≥29 °C) at 1.5 °C and 2.7 °C global warming: **a**. Absolute land area exposed, for the top 50 countries under 2.7 °C global warming (note the break in the x-axis). **b**. Fraction of land area exposed, for the top 50 countries under 2.7 °C global warming.

## Extended Data Tables

**Extended Data Table 1.** Downscaled CMIP6 climate model runs used in this study for different time intervals and SSP (‘S’) scenarios. The model data were downloaded from the WorldClim v2.0 database (https://www.worldclim.org/).

| GCM | 2020-2040 |  |  |  | 2040-2060 |  |  |  | 2060-2080 |  |  |  | 2080-2100 |  |  |  |
| --- | --- | --- | --- | --- | --- | --- | --- | --- | --- | --- | --- | --- | --- | --- | --- | --- |
|  | S1 | S2 | S3 | S5 | S1 | S2 | S3 | S5 | S1 | S2 | S3 | S5 | S1 | S2 | S3 | S5 |
| BCC-CSM2-MR | ✓ | ✓ | ✓ | ✓ | ✓ | ✓ | ✓ | ✓ | ✓ | ✓ | ✓ | ✓ | ✓ | ✓ | ✓ | ✓ |
| CNRM-CM6-1 | ✓ | ✓ | ✓ | ✓ | ✓ | ✓ | ✓ | ✓ | ✓ | ✓ | ✓ | ✓ | ✓ | ✓ | ✓ | ✓ |
| CNRM-ESM2-1 | ✓ | ✓ | ✓ | ✓ | ✓ | ✓ | ✓ | ✓ | ✓ | ✓ | ✓ | ✓ | ✓ | ✓ | ✓ | ✓ |
| CanESM5 | ✓ | ✓ | ✓ | ✓ | ✓ | ✓ | ✓ | ✓ | ✓ | ✓ | ✓ | ✓ | ✓ | ✓ | ✓ | ✓ |
| GFDL-ESM4 | ✓ |  | ✓ |  | ✓ |  | ✓ |  | ✓ |  | ✓ |  | ✓ |  | ✓ |  |
| IPSL-CM6A-LR | ✓ |  | ✓ | ✓ | ✓ | ✓ | ✓ | ✓ | ✓ | ✓ | ✓ | ✓ | ✓ | ✓ | ✓ | ✓ |
| MIROC-ES2L | ✓ | ✓ | ✓ | ✓ | ✓ | ✓ | ✓ | ✓ | ✓ | ✓ | ✓ | ✓ | ✓ | ✓ | ✓ | ✓ |
| MRI-ESM2-0 | ✓ | ✓ | ✓ |  | ✓ | ✓ | ✓ | ✓ | ✓ | ✓ | ✓ | ✓ | ✓ | ✓ | ✓ | ✓ |

**Extended Data Table 2.** Global warming (20-year averages) from the full CMIP6 ensemble (Table SPM.1 of IPCC AR6 WG1) and world population levels (central year) for each Shared Socioeconomic Pathway (SSP).

| Warming<br>(°C) | Scenario | 2020-2040 | 2040-2060 |  | 2080-2100 |
| --- | --- | --- | --- | --- | --- |
|  | SSP1-2.6 | 1.5 (1.2-1.8) | 1.7 (1.3-2.2) |  | 1.8 (1.3-2.4) |
|  | SSP2-4.5 | 1.5 (1.2-1.8) | 2.0 (1.6-2.5) |  | 2.7 (2.1-3.5) |
|  | SSP3-7.0 | 1.5 (1.2-1.8) | 2.1 (1.7-2.6) |  | 3.6 (2.8-4.6) |
|  | SSP5-8.5 | 1.6 (1.3-1.9) | 2.4 (1.9-3.0) |  | 4.4 (3.3-5.7) |
| World total<br>Population<br>(billion) | Scenario | 2030 | 2050 | 2070 | 2090 |
|  | SSP1 | 8.0 | 8.5 | 8.2 | 7.4 |
|  | SSP2 | 8.3 | 9.2 | 9.5 | 9.2 |
|  | SSP3 | 8.5 | 9.9 | 11.1 | 12.1 |
|  | SSP5 | 8.0 | 8.6 | 8.4 | 7.8 |

